# Population genomics of *Wolbachia* and mtDNA in *Drosophila simulans* from California

**DOI:** 10.1101/174375

**Authors:** Sarah Signor

**Affiliations:** Department of Molecular and Computational Biology, University of Southern California, Los Angeles, California USA

**Keywords:** *Drosophila simulans*, population genetics, *Wolbachia*

## Abstract

*Wolbachia pipientis* is an intracellular endosymbiont infecting many arthropods and filarial nematodes. Little is known about the short-term evolution of *Wolbachia* or its interaction with its host. *Wolbachia* is maternally inherited, resulting in co-inheritance of mitochondrial organelles such as mtDNA. Here I explore the short-term evolution of *Wolbachia*, and the relationship between *Wolbachia* and mtDNA, using a large inbred panel of *Drosophila simulans* infected with the *Wolbachia* strain *w*Ri. I find reduced diversity relative to expectation in both *Wolbachia* and mtDNA, but only mtDNA shows evidence of a recent selective sweep or population bottleneck. I estimate *Wolbachia* and mtDNA titre in each genotype, and I find considerable variation in both phenotypes, despite low genetic diversity in *Wolbachia* and mtDNA. A phylogeny of *Wolbachia* and of mtDNA show that both trees are largely unresolved, suggesting a recent origin of the infection and a single origin. Using Wolbachia and mtDNA titre as a phenotype, we perform an association analysis with the nuclear genome and find several regions implicated in the phenotype, including one which contains four CAAX-box protein processing genes. CAAX-box protein processing can be an important part of host-pathogen interactions in other systems, suggesting interesting directions for future research.

## INTRODUCTION

Heritable symbiotic associations such as that between *Drosophila* and *Wolbachia pipientis* have widespread impact on host ecology and evolution. These types of heritable endosymbiotic relationships are recognized as key drivers of evolution, but the intraspecific variation that effects their short-term evolution is not well explored. *Wolbachia* are α-proteobacterial endosymbionts found in up to 40% of all arthropod species^1-3^. *Wolbachia* are maternally transmitted and spread through manipulating the reproductive strategies of their host, using mechanisms such as feminization, male-killing, or cytoplasmic incompatibility. The most common of these is cytoplasmic incompatibility, where mating between males and females of the same species results in embryonic mortality if they have different *Wolbachia* infection status^4-8^. *Wolbachia* may also confer certain protections upon their host, such as increased resistance to certain viruses, or increased survival when exposed to certain environmental stressors^9-15^. *Wolbachia* is one of the most abundant obligate intracellular parasites, given that 85% of animal species are insects. This has profound meaning for evolutionary processes such as sexual selection and speciation^16,17^.

*Wolbachia* strain *w*Ri is known to have spread recently in the sister species to the model organism *Drosophila melanogaster*, *D. simulans*^4,5,8,18^. It was at ∼95-100% frequency in Southern California populations at the time its original sampling in the 1980’s^19,20^. It likely invaded California less than 25 years before it was first detected in 1984^21^. It is now thought to have been horizontally transmitted to *D. simulans* from *D. ananassae*, though the same strain is also found in *D. suzukkii*^22^. The maternal transmission of *Wolbachia* means that as the microorganism spreads all maternally inherited organelles spread along with it. Most notably mtDNA will be forced through a bottleneck, lowering the diversity of mtDNA in infected populations^18,23,24^. This will cause mtDNA and *Wolbachia* to be more closely associated than nuclear genes, and this coupling has been demonstrated previously in *D. simulans*^18,25,26^. In fact, *D. simulans* is known to have three major mitochondrial haplotypes (*si*I, *si*II, and *si*III) and two subtypes (*si*IIA and *si*IIB) that harbor very little variation and that appear to be nonrandomly associated with *Wolbachia* strains^27-29^. These mitochondrial haplotypes are largely allopatric, except for the presence of both *si*II, and *si*III in Madagascar and La Reunion^30^.

In *D. melanogaster* variation in *Wolbachia* has been well investigated using genomic data, though this has not been done in the genomic era in *D. simulans*^31^. They found long lived associations between mitochondrial and *Wolbachia* haplotypes and strong geographic structuring among cytotypes^31-33^. This study also observed that *Wolbachia* titre varied among fly populations as the result of intraspecific nuclear genetic variation^31^. However, the assumption that it was due to intraspecific nuclear background was based on the presence of a constant environment and no polymorphisms were identified that could be affecting this phenotype. Very little is known about how *Wolbachia* interact with their hosts, though recent work has uncovered evidence that deubiquitylating enzymes produced by *Wolbachia* and secreted into the host cytoplasm mediate cytoplasmic incompatibility^34^. *Wolbachia* DNA is also frequently inserted into the host genome, though this has not occurred with *w*Ri in *D. simulans*^21^. Genes involved in the formation of germline stem cells such as *benign gonial cell neoplasm* and *bag-of-marbles* are considered candidates for interacting with *Wolbachia*, and have been found to have unusual population genetic patterns in *D. melanogaster* ^35,36^. *bag-of-marbles* has been suggested to interact with *Wolbachia* due to fertility rescue in hypomorphs, but the interaction of this gene with *Wolbachia* in natural populations is not clear^22,35-37^. Notably, *Wolbachia* localizes in tissues differently depending upon the strain and species so the interactions between the host and *Wolbachia* are likely to also be different^38,39^.

*Wolbachia* infections must be maintained in host populations through transovarial transmission, wherein *Wolbachia* is present in the germline at sufficient copy number to ensure transmission but not to cause host pathology^40^. *Wolbachia* titre has been shown to have important phenotypic effects on the host^11,41-49^. However, control of *Wolbachia* replication is not well understood, nor is the dependence of this control on host background versus bacterial genotype^11,50-52^. Differences in *Wolbachia* titre when it is transinfected between species suggests a role of host background in controlling copy number, population genomics in *D. melanogaster* suggest an effect of host background, and there does seem to be host-specific patterns of tissue colonization^53-55^. However, multiple *Wolbachia* genotypes can also behave differently in the same genetic background suggesting contributions from the bacterial genome^51,56^. It is also possible to select for greater *Wolbachia* densities, though the heritability of this is unclear^57,58^.

Here I investigate the dynamics of *Wolbachia* and mtDNA in a large panel of *D. simulans* from a single Californian population. I determine infection status of *Wolbachia* in the panel of *D. simulans* genotypes. I look for signatures of selection in both genomes using summary statistics Tajima’s *D* and *π* and find that while *Wolbachia* patterns of variation are not unusual given its demographic history the reduction in mtDNA diversity is suggestive of a recent bottleneck due either to selection or changes in population size. I also measure linkage disequilibrium between mtDNA and *Wolbachia* as a proxy for coinheritance. Using whole genome sequences, I investigate the phylogeny of both *Wolbachia* and mtDNA and find that in this population they are essentially unresolved. I investigate variation in the copy number of both *Wolbachia* and mtDNA in this population using relative estimates derived from illumina sequencing coverage compared to nuclear coverage. I find considerable copy number variation in this population, and an association analysis using this as a phenotype implicates several genomic regions potentially involved in mediating this phenotype. This includes a region containing multiple genes invovled CAAX-box protein prenylation, a process that is important for mediating the relationship between host and pathogen in other systems^59-61^.

## METHODS

### *Drosophila* strains

Strains are as described in^62^. Briefly, the 167 *D. simulans* lines were collected in the Zuma organic orchard in Zuma beach, California in February of 2012 from a single pile of fermenting strawberries. Single mated females were collected and inbred by 15 generations of full sib mating of their progeny. *Drosophila* were raised at a constant temperature of 20º C with 12-hour light/dark cycles. They were raised on a standard glucose/yeast media, and each library was constructed from adult females of similar age (less than one week).

### Data sources and processing

The sequencing reads were downloaded from the NCBI Short Read Archive from project SRP075682. Libraries were assembled using BWA mem (v. 0.7.5), and processed with samtools (v. 0.1.19) using default parameters^63,64^. The *Wolbachia* reference is the *w*Ri strain previously identified in Southern California (Accession number NC_012416)^21^. The mtDNA reference is from *D. simulans w*^*501*^, which is haplogroup *siII* as expected for *D. simulans* from California (Accession number KC244284)^65^. PCR duplicates were removed using Picard MarkDuplicates (v. 2.9.4) and GATK (v. 3.7) was used for indel realignment and SNP calling using default parameters (http://picard.sourceforge.net)^66^. SNPs were called jointly for all genotypes using Haplotypecaller^66^. Individual consensus fasta sequences were produced using SelectVariants to create individual vcf files and FastaAlternateReferenceMaker. Vcf files were filtered for indels and non-biallelic SNPs using VCFtools (v. 0.1.13)^67^. The files were also filtered for SNPs with more than 10% missing data. The *Wolbachia* genome was filtered for regions of unusual coverage or SNP density, for example two regions of the *Wolbachia* genome harbored ∼40 SNPs within two kb, far above background levels of variation (Supp. Fig.1). These two regions coincided with regions of unusually high coverage suggesting they are repeated elements.

### Prediction of *Wolbachia* infection status

*Wolbachia* infection status was determined by calculating the mean depth of coverage of the assembly and the breadth of coverage of the consensus sequence using bedtools^68^. Depth of coverage refers to the average read depth across the Wolbachia genome, while breadth of coverage refers to the number of bases covered by at least two reads. Depth of coverage at each nucleotide was estimated using the genomecov function, while breadth was estimated using the coverage function. Predictions of *Wolbachia* infection status using illumina data have previously been shown to have 98.8% concordance with PCR based predication of infection status^32^.

### Nucleotide diversity

Levels of polymorphism for mtDNA and *Wolbachia* were estimated as *π* in 10 kb windows using VCFtools (v0.1.14)^69^. To investigate whether the frequency spectrum conformed to the standard neutral model of molecular evolution I also calculated Tajima’s *D* in 10 kb windows using VCFtools. To assess the significance of deviations in Tajima’s *D* and *π* 10,000 simulations were performed using msms conditioned on the number of variable sites and with no recombination^70^.

### Linkage disequilibrium

Linkage between *Wolbachia* and mtDNA SNPs could potentially be a predictor of co-inheritance of mtDNA and *Wolbachia*. Linkage was estimated using VCFtools (v0.1.14) using inter-chrom-geno-r2 to estimate r^2^ between ach SNP in the two genomes^67^.

### Estimation of mtDNA and *Wolbachia* copy number

In insects, the phenotypic effect of *Wolbachia* will vary depending upon copy number in the host cells^9,32^. Given that there are two copies of autosomal DNA in a cell, I infer mtDNA and *Wolbachia* copy number based on the ratio between mtDNA and autosomal DNA. This is intended to provide a relative estimate of copy number rather than an absolute measure. Relative copy number estimated in this way obscures intra-individual variation and variation between tissues, though the authors note that all flies used in constructing the libraries were females of approximately the same age. *Wolbachia* contains several regions which were excluded due to unusually high coverage across all samples (more than 3x the mean coverage). Average coverage of autosomal DNA was calculated from randomly chosen and equivalently sized nuclear regions for each mtDNA (Scf_3L:8000000..8014945) and *Wolbachia* (Scf_2L:11000000..11445873). The average coverage of each nuclear region, respectively, was then used to normalize estimates of copy number for each genotype. Previously the results of measuring *Wolbachia* copy number in the same samples using both qPCR and estimates from illumina read depth had a Pearson’s correlation coefficient of .79, thus this is a robust approach to measuring *Wolbachia* titre^31^.

### Phylogenomic analysis

To understand the relationship between *Wolbachia* infection and mtDNA I reconstructed the genealogical history of each within the sample population. Multiple alignments were generated for both mtDNA and *Wolbachia* by concatenating fasta consensus sequence files for each genotype. All indels and non-biallelic SNPs were excluded from the dataset prior to generating the consensus fasta for each genotype. RAxML version 8.10.2 was used to reconstruct phylogenies^71^. Maximum likelihood tree searches were conducted using a general time reversible (GTR) model of nucleotide substitution with CAT rate heterogeneity and all model parameters estimated by RAxM^72^. Trees were inferred using the rapid bootstrap algorithm and simultaneous estimation of trees and bootstrapping, with automatic estimation of the necessary number of bootstrap replicates.

### Association Analysis

The association analysis focused on a relationship between nuclear polymorphisms and *Wolbachia* and mtDNA copy number. To reduce the need for correction due to multiple testing and focus on regions that may have been affected by selection due to the recent invasion of *Wolbachia* I used a subset of the genome identified previously as exhibiting haplotype structure suggestive of recent selection^62,73^. These regions are unusually long haplotype blocks, thus many of the SNPs within each block are not independent, reducing the need for correction due to multiple testing. Heterozygous bases were coded as missing, and all loci with more than 10% missing data were excluded from the analysis, as well as SNPs with a minor allele frequency of less than 2%, meaning they were present in the population in at least 3 copies. mtDNA and *Wolbachia* copy number were used for a multivariate analysis of association using plink.multivariate^74^. To investigate the possibility that *Wolbachia* copy number is affected by polymorphisms in mtDNA, and vice versa, a single trait analysis was performed using plink v. 1.07^75^.

## RESULTS

### Sequencing Data

The autosomal data included in this analysis was reported in^62^. There was very little variation in both *Wolbachia* and mtDNA in this population. This included 78 SNPs and indels in the *Wolbachia* genome and 90 in mtDNA. Reduced diversity has been reported previously in *D. simulans* mtDNA^24,25^. The authors note that previous work has established that there is no unusual relatedness in the nuclear genome of this population^62^.

### Infection status

In a previous study lines were scored as infected if they had a breadth of coverage greater than 90% and a mean depth greater than one^32^. However, that dataset had a clearly bimodal distribution between infected and uninfected lines, where uninfected lines had breadth of coverage less than 10% while infected lines had a breadth of coverage of greater than 90%. As such that this demarcation was a natural interpretation of the data^32^. In *D. simulans*, all lines had ∼99% breadth of coverage aside from a single line with both a lower overall depth of coverage and 80% breadth (Fig. 1). For this reason, all lines were scored as infected. 100% infection is not unusually high for *D. simulans*.

**Figure 1.**
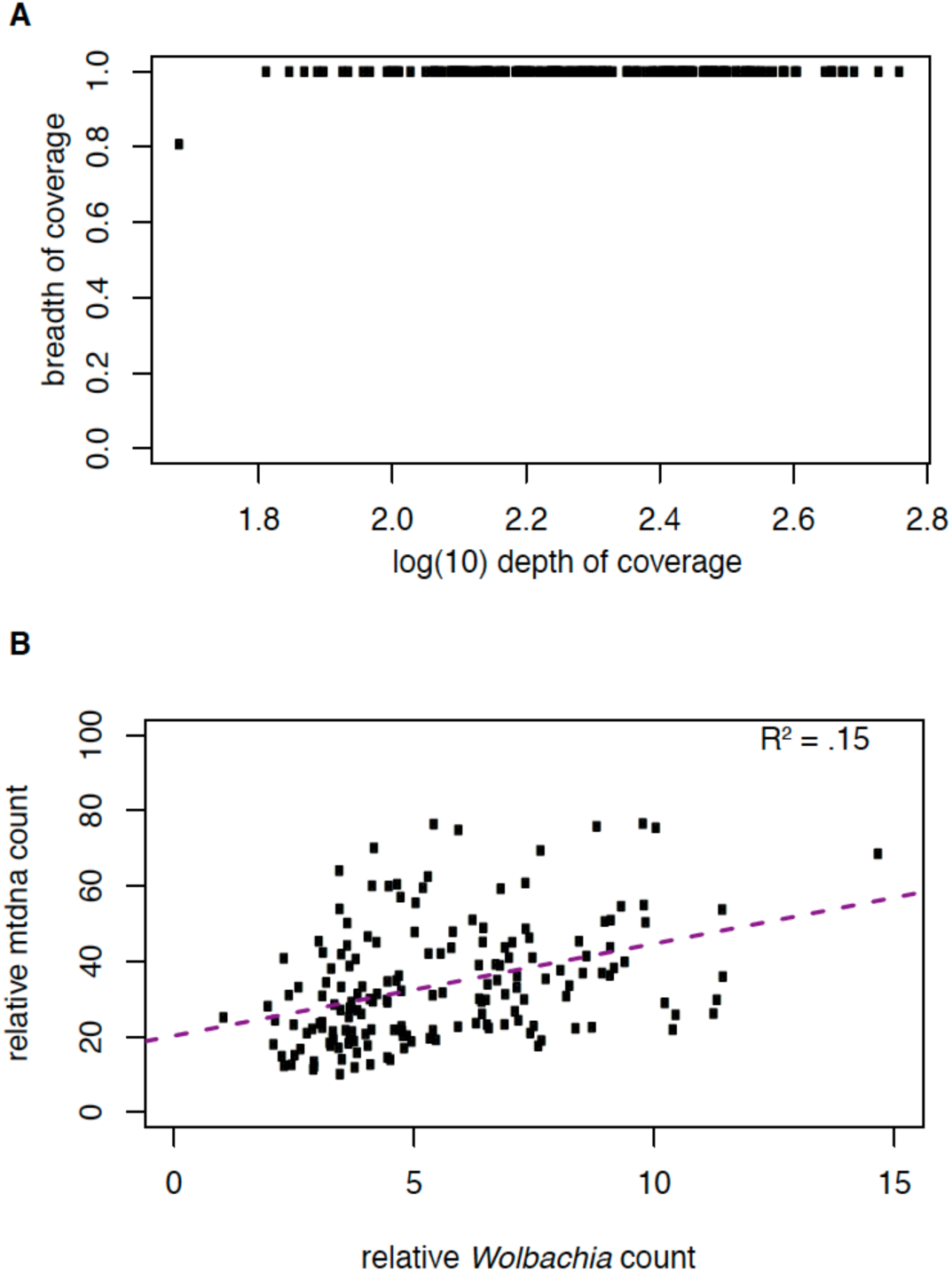
*Wolbachia* infection status and relationship to mtDNA copy number A. Relationship between depth and breadth of sequencing coverage for *Wolbachia* assemblies in the *D. simulans* panel. Depth of coverage is shown in log_10_ unites and is calculated as the number of reads present at each nucleotide in the reference averaged over every site. Breadth of coverage is the proportion of covered nucleotides in the consensus sequence relative to the reference. **B.** Relationship between relative mtDNA copy number and *Wolbachia* copy number. Both were normalized relative to nuclear coverage. Although separate regions were used to normalize mtDNA and *Wolbachia*, as they are different sizes, average values were very similar within genotypes. The relationship between mtDNA and *Wolbachia* copy number is positive (p < 2.4 x 10^-7^).

### Nucleotide Diversity

Estimates of *π* in *Wolbachia* ranged from 5.98 x 10^-7^ to 1 x 10^-3^, with an average of 1.42 x 10^-5^, within the range of estimates from *Wolbachia* in *D. melanogaster* from another study (7.9 x 10^-6^ – 2.8 x 10^-5^)^31^. The mean of *π* in simulated populations of *Wolbachia* is 1.9 x 10^-3^ suggesting that variation is somewhat reduced in *w*Ri. *π* in mtDNA is 1 x 10^-4^ which again is similar to estimates from *D. melanogaster* (4.34 x 10^-4^ – 1.51 x 10^-3^)^31^.

Overall Tajima’s *D* was estimated to be -2.4 for *D. simulans* mtDNA (Fig 2). This is similar to estimates in *D. melanogaster*^32^. Significance of this estimate was assessed using 10,000 simulations in msms conditioned on the number of segregating sites and no recombination, and it is significant at *p* < .05. Tajima’s *D* in *Wolbachia* is not significantly different from expectations under neutrality based on 10,000 simulations. Thus, while a selective sweep or population bottleneck seems to have strongly effected mtDNA in *D. simulans*, the same is not true of the *Wolbachia* population (Fig 2). This is very different from *D. melanogaster* where *Wolbachia* and mtDNA had similar patterns of nucleotide diversity^32^.

**Figure 2.**
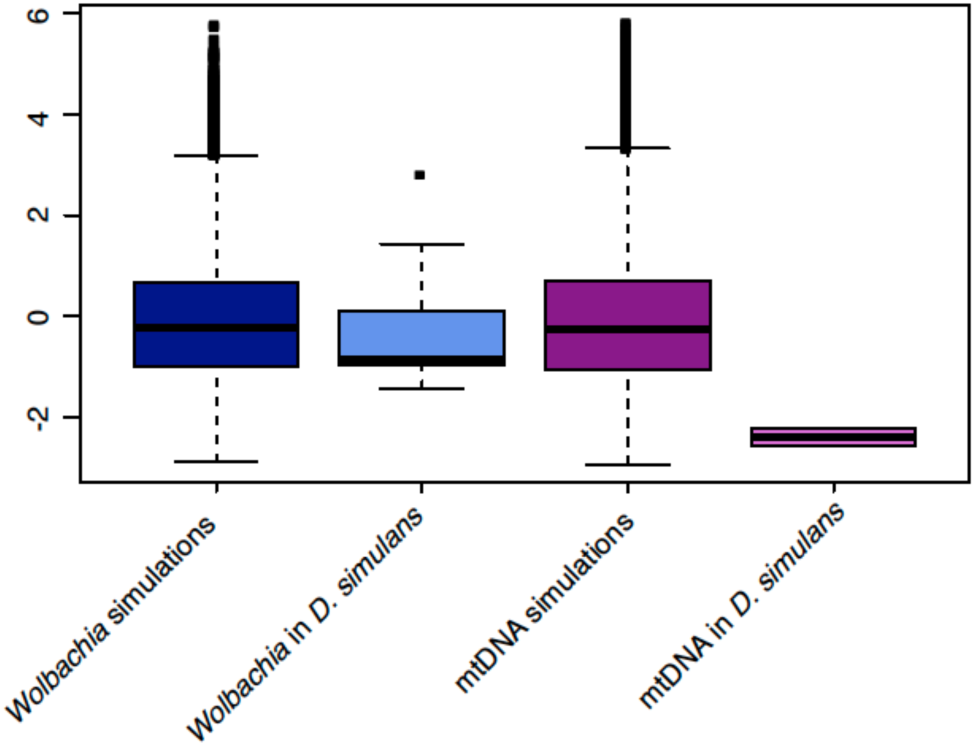
*Wolbachia* and mtDNA Tajima’s *D*. 10,000 simulations were performed for *Wolbachia* and *D. simulans* each conditioned upon the number of polymorphisms. The actual values in *D. simulans* mtDNA are outside the 95% confidence interval of the simulations, while *Wolbachia* is not. There is considerable variation in Tajima’s *D* across the *Wolbachia* genome while mtDNA is much smaller and invariant in its values of Tajima’s *D*.

This is also much more negative than previously reported for mtDNA in *D. simulans*^25^. It is very different from the general patterns of Tajima’s *D* in the nuclear genome, where average Tajima’s *D* is 1 and the majority of the genome has a positive Tajima’s *D*. Simulations in previous work suggest that the pervasively positive values in the nuclear genome may be due to a population contraction, which again indicates that the population dynamics affecting *D. simulans* nuclear and mtDNA genomes are very different^25,62^.

### Linkage disequilibrium

There was no significant linkage disequilibrium between the genomes of *Wolbachia* and *D. simulans* mtDNA. Average LD between *Wolbachia* and mtDNA SNPs was 2.06 x 10^-3^. This may be because the infection of *D. simulans* was too recent for variation to accumulate along particular lineages, and also suggests that *D. simulans* was infected by a single invasion.

### Estimation of mtDNA and *Wolbachia* copy number

There was considerable heterogeneity in both *Wolbachia* and mtDNA copy number (Fig. 1). Mean (standard deviation) copy number of *Wolbachia* is 5.56 (2.45). This is similar to one estimate in *D. melanogaster*, where mean copy number is 5.57 (3.95) though the standard deviation is lower in *D. simulans* ^32^. The reported mean was lower in other populations of *D. melanogaster*, though still within the same range (2 - 4.5)^31^. Similarly mean mtDNA copy number is 33.85 (15.5) in *D. simulans* and 32.9 (44.5) in one estimate for *D. melanogaster*^32^. This is again not an absolute measure, but relative to nuclear genomic coverage. The lower standard deviation could be due to more precise staging of the age of *D. simulans*, less background variation effecting copy number (the *D. melanogaster* sample was from multiple populations), or other unknown mechanisms. There was a positive relationship between mtDNA and *Wolbachia* copy number (Fig. 1) (*p* < 2.4 x 10^-7^). While the functional reasons for or consequences of this are unclear, because they are correlated they will be used in a multivariate analysis of association rather than as separate analyses.

### Phylogenomic analysis

To understand the relationship between *Wolbachia* infection status and mtDNA sequence variation I reconstructed the phylogenetic history of the complete *Wolbachia* and mtDNA genome using the entire set of 167 strains (Fig 3-4). What I found is consistent with the recent spread of *Wolbachia* in *D. simulans*, as both phylogenies are essentially unresolved. This is not unexpected for mtDNA given previous work in the species which found little within-haplotype variation among the three major mtDNA haplotypes in *D. simulans*^25,28^. Furthermore, of the 167 sequences 88 are identical to at least one other sequence in the sample. While the *Wolbachia* phylogenetic tree gives the impression of having more resolution than mtDNA, this is likely due to the larger genome, as the branches have similarly low support. Of the 167 strains included in the tree 18 are identical to one or more *Wolbachia* genomes. Both trees are essentially star phylogenies with the majority of bootstrap support values being less than 30. Bootstrap support of greater than 70, for two branches in the mtDNA tree and five in the *Wolbachia* tree, is shown (Fig 3-4). If uninfected individuals had been included in the dataset perhaps it would be possible to test for congruence between the two phylogenies, however the essentially unresolved trees make it clear that both *Wolbachia* and mtDNA swept the population recently.

**Figure 3:**
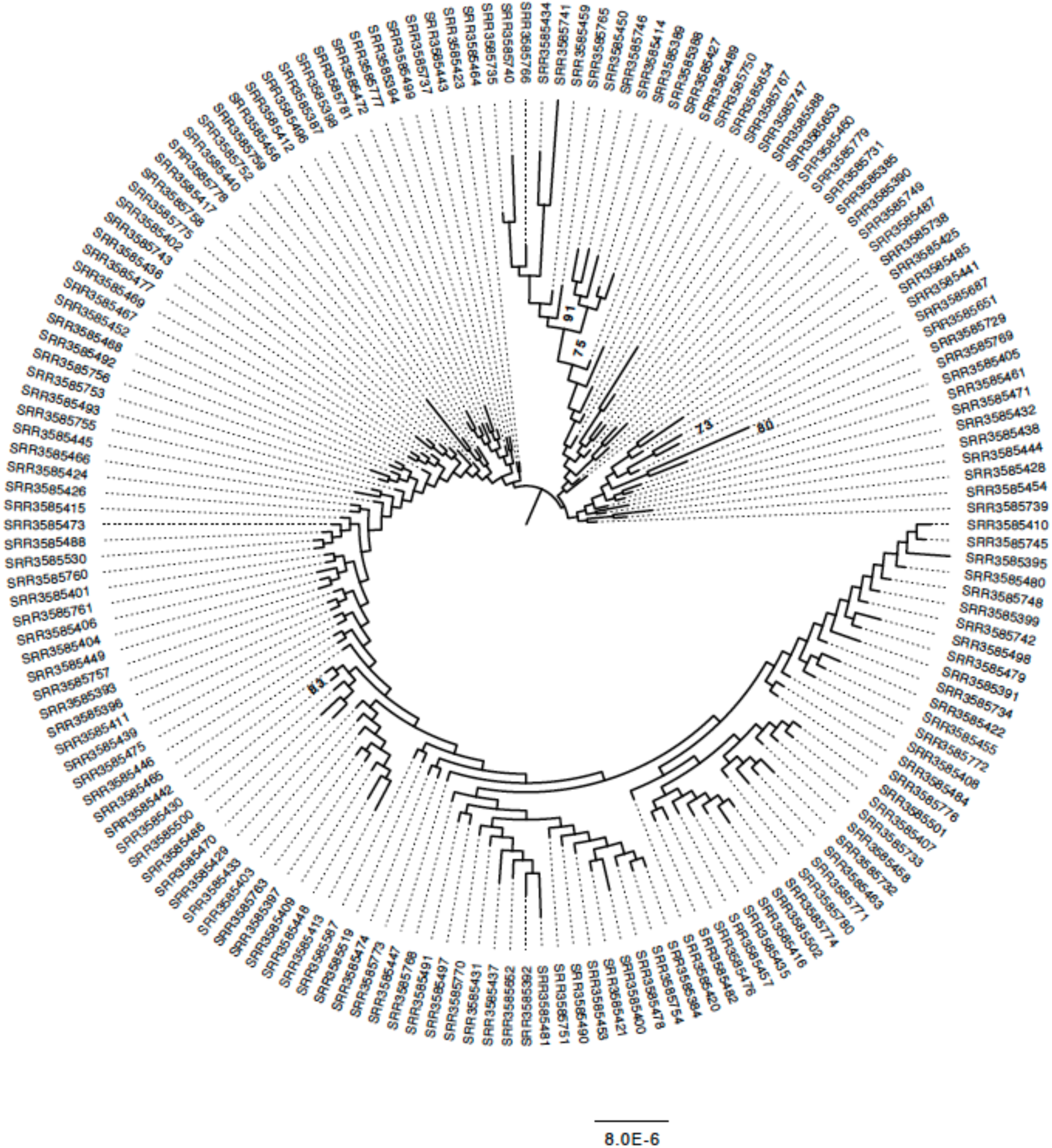
Maximum likelihood genealogy of the *D. simulans Wolbachia* pathogen. All strains were infected with *Wolbachia* and are included in this geneaology. The underlying data consist of an ungapped multiple alignment of 168 sequences of the entire *Wolbachia* genome. The unrooted tree was midpoint rooted for visualization and branches with > 70% RAxML bootstrap support values are shown in bold. Scale bars for branch lengths are in term of mutations per site. The majority of branches are essentially unsupported by bootstrapping.

**Figure 4:**
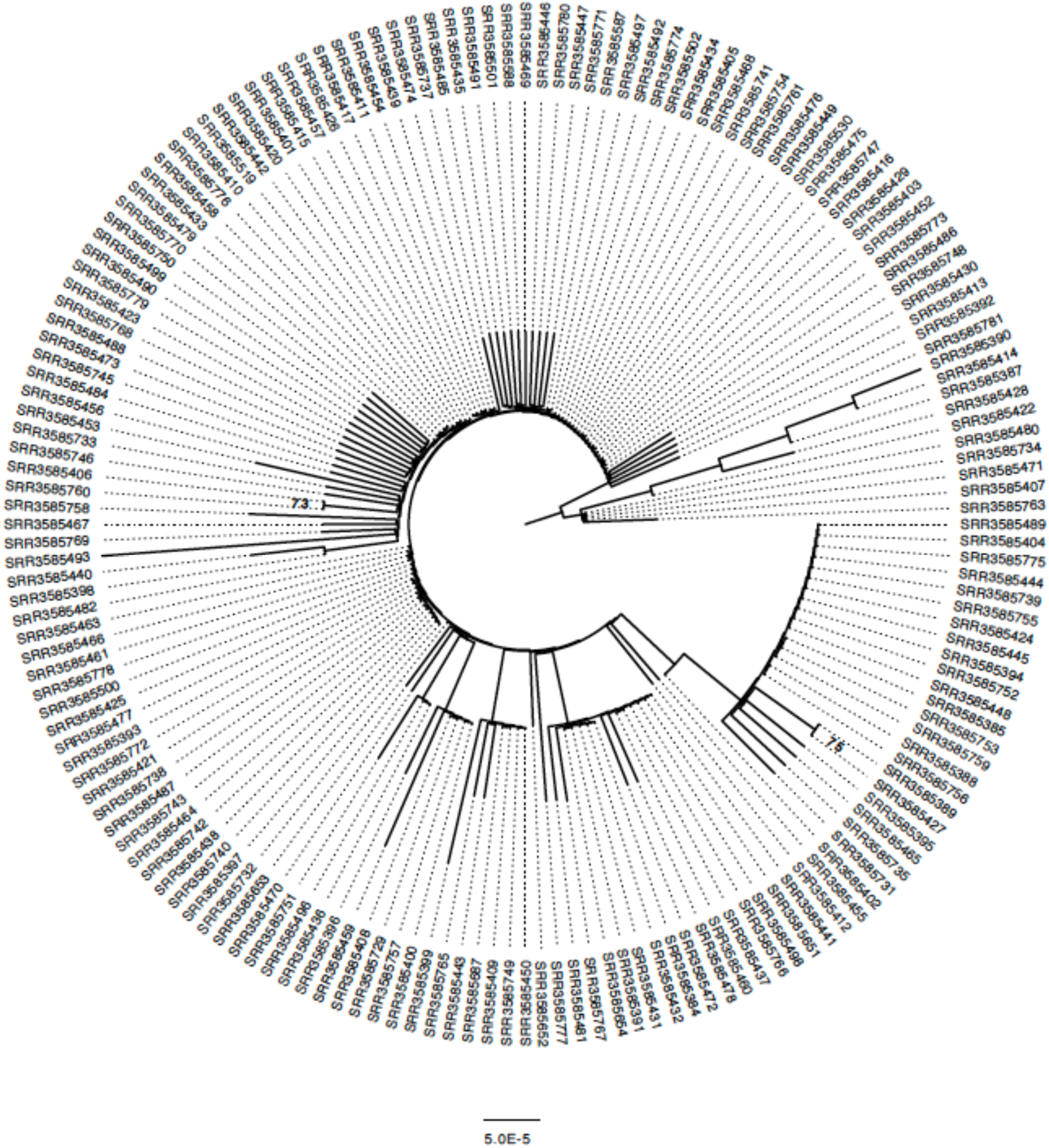
Maximum likelihood genealogy of the *D. simulans* mtDNA genome. The underlying data consist of an ungapped multiple alignment of 168 sequences of the entire mtDNA genome. The unrooted tree was midpoint rooted for visualization and branches with > 70% RAxML bootstrap support values are shown in bold. Scale bars for branch lengths are in term of mutations per site. The tree is largely unresolved, suggesting recent spread of this mtDNA haplotype through the population.

### Association Analysis

Association analysis was performed using plink.multivariate by regressing the line means for mtDNA and *Wolbachia* copy number on each SNP contained within the previously identified in a scan for selection^62^. This scan for selection focused on identifying haplotype blocks in LD. This considerably reduces the number of SNPs tested for association, in addition to the fact that the SNPs are in haplotype blocks and are therefore not independent tests^62,73^. This reduces the need for correction due to multiple testing. I used a *p*-value cut-off of *p* < 9 x 10^-6^ and identified 16 SNPs associated with *Wolbachia* and mtDNA copy number. Of these 16 SNPs 13 are located in the same region on chromosome 2R (Scf_2R: 13550916-13569038). Given the concentration of significant SNPs in a single region, this is also the region we will focus on the most in the following discussion. The region containing 13 SNPs contains nine genes, four of which are involved in CAAX-box protein processing, *ste24a-c* and a recent duplicate of *ste24c* CG30461. CAAX-box protein processing is a part of a series of posttranslational protein modifications collectively called protein prenylation which are required for fully functional proteins to be targeted to cell membranes or organelles. It has been shown that pathogenic bacteria can exploit the host cell’s prenylation machinery, though it is unclear if this occurs in *Wolbachia*^59^.

The other five genes are *AsnRs-m*, which is largely unannotated but is thought to a mitochondrial aminoacyl-tRNA synthetase^76^. *NIPP1Dm* is involved in axon guidance and negative regulation of protein phosphorylation^77,78^. *CG6805* is generally unannotated but is inferred to be involved in dephosporylation^76^. *Cbp53E* regulates neural development^79^. Lastly, *Ehbp1* is a developmental gene implicated in regulation of the Notch pathway and membrane organization^80^.

Of the other three SNPs identified in this association analysis two are located at Scf_2R:5814103 and Scf_2R:5811043, while the third is located at Scf_3L:2055556. Scf_2R:5811043 and Scf_2R:5814103 are located in *Su(var)2-10* and *Phax*, respectively. These are neighboring genes, though there is a third gene within 10 kb, *Mys45A*. *Su(var)2-10* is involved in development and chromosome organization, but it has also been implicated in the regulation of the innate immune response and defense against gram-negative bacteria^81^. *Su(var)2-10* is of particular interest given that *Wolbachia* are gram-negative bacteria, however the potential role of *Su(var)2-10* in immune response is not clear. *Phax* is not well annotated but is inferred to be involved in snRNA export from the nucleus^79^. *Mys45A* is potentially involved in actin cytoskeleton organization^79^. In *D. melanogaster Wolbachia* uses host actin for maternal transmission, though this has not been verified in *D. simulans*^82^. The last SNP, at Scf_3L:2055556, is in *Connectin*, a cell adhesion protein also involved in axon guidance^83^.

The identification of these SNPs in association with mtDNA and *Wolbachia* copy number does not imply a functional relationship. Nonetheless, I chose to investigate whether any of these substitutions had an effect on the coding sequence of any of genes in the region. Of the three SNPs found outside the region containing the CAAX-box proteins all were either in introns or regulatory regions. Of the 13 SNPs identified between Scf_2R: 13550916-13569038 eight are in introns or untranslated regions, including one in the long intron of *Cb53E*, three in the introns or noncoding transcript of *CG6805*, and two in the introns of *Ephb*. Of the remaining five SNPs four are in coding regions but silent, causing no change in the amino acid sequence of the protein. This includes silent mutations in the exons of *ste24c* and two silent mutations in the exons of *Epbh*. One SNP located in an exon of *ste24a*, at 13558515, is an amino acid substitution from a Leucine to a Valine. This is not an uncommon amino acid substitution^84,85^, though it can be associated with phenotypes^86,87^. Mutations in introns and untranslated regions could also be having an effect on gene expression or processing, as could other linked SNPs in the region that were not included in the analysis.

*Association between Wolbachia and mtDNA*

Association analysis was performed using plink by regressing the line means for mtDNA copy number onto the *Wolbachia* genome and vice versa^75^. There was no association between *Wolbachia* SNPs and mtDNA copy number, but the opposite was not true. One SNP in the *D. simulans* mtDNA affected *Wolbachia* copy number at *p* < 3.18 x 10^-6^. It is located in the *D. simulans* homolog of *D. melanogaster srRNA* which has been implicated in pole cell formation^88^. *Wolbachia* is incorporated into the pole cells, the precursor to the germline, in order to be transmitted^89^.

## DISCUSSION

Using high through-put sequencing of a large panel of *D. simulans* I have reconstructed the complete genome sequences of mtDNA and *Wolbachia*. I use these genome sequences to investigate the recent history of *Wolbachia* and mtDNA in this population, as well as to estimate titre of both *Wolbachia* and mtDNA. The history of *Wolbachia* in this population is reflected in the essentially star-like phylogeny of both mtDNA and *Wolbachia*, indicating recent spread and co-inheritance. Lack of variation at mtDNA and *Wolbachia* suggests a single spread of *w*Ri in this population as well as strict vertical transmission in the maternal cytoplasm. Variation in *Wolbachia* is within the range expected under a neutral model, however that was not the case for mtDNA which suggests either a selection sweep or a population bottleneck. Previous studies found similar population genetic patterns at *Wolbachia* and mtDNA in *D. melanogaster*, and thus could not distinguish whether selection on *Wolbachia* was driving similar patterns in mtDNA or vice versa^31^. The much stronger pattern of negative Tajima’s *D* in the mtDNA suggests that in *D. simulans* selection is in fact mitochondrial. There was no linkage disequilibrium between *Wolbachia* and mtDNA variants, however this is most likely due to fixation of a single mitochondrial haplotype without considerable subsequent mutation.

Currently little is known about how *Wolbachia* interacts with its host^37-39,82,90^. Understanding these interactions, including regulation of *Wolbachia* titre, will be key to understanding the evolution of *Wolbachia* and its hosts. By normalizing *Wolbachia* and mtDNA copy number using coverage of the nuclear genome I am able to obtain estimates of its abundance. Much as in previous work, mtDNA copy number was higher than *Wolbachia* copy number, though both varied across strains^32^. As all of my data was produced from adult females, at the same time, using the same techniques, there is no danger that this is due to differences in methodology among samples^32^. Estimates of copy number were very similar to previous work in *D. melanogaster*, performed with qPCR, and there has been shown to be a high correlation between qPCR and illumina estimates of copy number^31,32^. These are not absolute measures, rather they are relative to one another and to nuclear copy number, and they provide robust estimates of *Wolbachia* titre within the population. As the *Wolbachia* phylogenetic tree is essentially unresolved in this population but there is considerable variation in *Wolbachia* titre, it is clear that some host factors must be affecting variation in *Wolbachia* titre.

The history of mtDNA and the nuclear genome is quite divergent in this population. The nuclear genome has an average Tajima’s *D* of 1 and 5 polymorphisms for every 100 bp^62^. Simulations suggest that this is due to a combination of population contraction and selection, most likely from standing variation, though many types of sweeps can produce similar signatures^62^. In contrast the mtDNA genome contains an abundance of low frequency variation, and in fact many of the majority of mtDNA genomes sampled in this population are identical. This is consistent with the recent spread, single origin, and maternal transmission, of *w*Ri in *D. simulans*. This is consistent with previous work which found low levels of mtDNA variation in *D. simulans* within a haplotype^24,91^. This is also consistent with work on *Wolbachia* which documented the spread of *w*Ri in *D. simulans* in the 1980’s^4,5,8,18-20^.

While it has been proposed elsewhere, the author is not aware of another association analysis of *Wolbachia* and mtDNA copy number^32^. *Wolbachia* copy number is known to be affected by host background, but the genes or mechanisms involved are not known^54,55,57^. The fact that four of the nine genes found in the primary region detected in the association analysis are involved in CAAX-box protein processing is of particular interest, given the history of this type of gene and intracellular pathogens. CAAX-box protein processing is a part of a series of posttranslational protein modifications collectively called protein prenylation which are required for fully functional proteins to be targeted to cell membranes or organelles. Prenylated proteins include Ras, Rac, and Rho. However, it has been shown that pathogenic bacteria can exploit the host cell’s prenylation machinery^59^. For example, *Salmonella-induced filament A* is a protein from *Salmonella typhimurium*, a gram-negative facultative intracellular bacterium. *Salmonella-induced filament A* has a CAAX motif required for prenylation to occur, it was shown to be processed by host prenylation machinery, and it is necessary for survival of the bacterium^60,92,93^. *Legionella pneumophila* Ankyrin B protein exploits the host prenylation machinery in order to anchor Ankyrin B protein to the membrane of the pathogenic vacuole^61^. Proliferation of *Legionella pneumophila* requires Ankyrin B, as does the manifestation of Legionnaires disease. Ankyrin repeat domains are most commonly found in eukaryotes and viruses, though they are rarely found in bacteria and Archaea^94^. In bacteria they are found in a few obligate or facultative intracellular Proteobacteria^59^. *Wolbachia* has an unusually high number of Ankyrin repeat domains with rapid evolution^94^. Ankyrin proteins play a major role in host-pathogen interactions and the evolution of infections^95,96^. There is no way to know from the current analysis if the Ankyrin repeat genes are exploiting the host prenylation system but it is an intriguing area for future investigation. The results of this association analysis suggest that some interaction between the pathogen and its host is targeting the protein prenylation machinery.

There was also an association between a polymorphism in *srRNA,* which has been implicated in pole cell formation^88^, and *Wolbachia* copy number. Concentration of *Wolbachia* in the posterior of the embryo, where pole cells are forming, is correlated with degree of cytoplasmic incompatibility^97^. *D. simulans* has been shown to have nearly complete cytoplasmic incompatibility, though it is possible there are mutations sorting at low frequency that affect this or that mitigate negative phenotypic consequences of high *Wolbachia* titre. It has also been demonstrated that *gurken* is important for *Wolbachia* titre in the germline in *D. melanogaster*, and it is involved in pole cell formation beginning at an earlier stage than *sr*RNA suggesting there could be an interaction between the two factors^88,90^. *D. simulans w*Ri has a different distribution in the cytoplasm from other strains of *Wolbachia*, as it tends to evenly distribute throughout the embryo while other strains are either concentrated at the posterior, or at the anterior of the embryo away from the pole cells^97^. Future work in related species may show that these different distributions also mitigate different interactions between host and symbiont, including being effected by different genes and processes within the host.

## Acknowledgements

I would like to thank J. Butler for helpful commentary on this manuscript, and S. Nuzhdin for helpful discussion and direction.

### Author Contributions

S. S. performed all of the work described herein.

### Competing financial interest

The author has no competing financial interests.

## References

1. Zug, R. & Hammerstein, P. Still a host of hosts for Wolbachia: analysis of recent data suggests that 40% of terrestrial arthropod species are infected. PLoS ONE 7, e38544 (2012).

2. Hilgenboecker, K., Hammerstein, P., Schlattmann, P., Telschow, A. & Werren, J. H. How many species are infected with Wolbachia-A statistical analysis of current data. FEMS Microbiol. Lett. 281, 215–220 (2008).

3. Jeyaprakash, A. & Hoy, M. A. Long PCR improves Wolbachia DNA amplification: wsp sequences found in 76% of sixty-three arthropod species. Insect Mol. Biol. 9, 393–405 (2000).

4. Turelli, M. & Hoffmann, A. A. Cytoplasmic incompatibility in *Drosophila simulans*: dynamics and parameter estimates from natural populations. Genetics 140, 1319–1338 (1995).

5. Turelli, M., Hoffmann, A. A. & McKechnie, S. W. Dynamics of cytoplasmic incompatibility and mtDNA variation in natural *Drosophila simulans* populations. Genetics 132, 713–723 (1992).

6. Charlat, S., Nirgianaki, A., Bourtzis, K. & Merçot, H. Evolution of Wolbachia-induced cytoplasmic incompatibility in *Drosophila simulans* and *D. sechellia*. Evolution 56, 1735–1742 (2002).

7. Rousset, F., Vautrin, D. & Solignac, M. Molecular identification of Wolbachia, the agent of cytoplasmic incompatibility in *Drosophila simulans*, and variability in relation with host mitochondrial types. Proc. R. Soc. B 247, 163–168 (1992).

8. Hoffmann, A. A., Turelli, M. & Harshman, L. G. Factors affecting the distribution of cytoplasmic incompatibility in *Drosophila simulans*. Genetics 126, 933–948 (1990).

9. Berticat, C., Rousset, F., Raymond, M., Berthomieu, A. & Weill, M. High Wolbachia density in insecticide-resistant mosquitoes. Proc. R. Soc. B 269, 1413–1416 (2002).

10. Wong, Z. S., Hedges, L. M., Brownlie, J. C. & Johnson, K. N. Wolbachia-mediated antibacterial protection and immune gene regulation in *Drosophila*. PLoS ONE 6, e25430–9 (2011).

11. Osborne, S. E., Leong, Y. S., O'Neill, S. L. & Johnson, K. N. Variation in antiviral protection mediated by different Wolbachia strains in *Drosophila simulans*. PLoS Pathog 5, e1000656–9 (2009).

12. Fytrou, A., Schofield, P. G., Kraaijeveld, A. R. & Hubbard, S. F. Wolbachia infection suppresses both host defence and parasitoid counter-defence. Proc. R. Soc. B 273, 791–796 (2006).

13. Brownlie, J. C. et al. Evidence for metabolic provisioning by a common invertebrate endosymbiont, *Wolbachia pipientis*, during periods of nutritional stress. PLoS Pathog 5, e1000368 (2009).

14. Teixeira, L., Ferreira, Á. & Ashburner, M. The bacterial symbiont Wolbachia induces resistance to RNA viral infections in *Drosophila melanogaster*. PLoS Biol. 6, e1000002 (2008).

15. Hedges, L. M., Brownlie, J. C. & O'Neill, S. L. Wolbachia and virus protection in insects.322, 702 Science (2008).

16. Jiggins, F. M., Hurst, G. & Majerus, M. Sex-ratio-distorting Wolbachia causes sex-role reversal in its butterfly host B. Proc. R. Soc. B 267, 69–73.

17. Koukou, K. et al. Influence of antibiotic treatment and Wolbachia curing on sexual isolation among *Drosophila melanogaster* cage populations. Evolution 60, 87–11 (2006).

18. Turelli, M. & Hoffmann, A. A. Rapid spread of an inherited incompatibility factor in California *Drosophila*. Nature 353, 440–442 (1991).

19. Hoffmann, A. A. & Turelli, M. Unidirectional incompatibility in *Drosophila simulans*: inheritance, geographic variation and fitness effects. Genetics 119, 435–444 (1988).

20. Hoffmann, A. A., Turelli, M. & Simmons, G. M. Unidirectional incompatibility between populations of *Drosophila simulans*. Evolution 40, 692–701 (1986).

21. Klasson, L. et al. The mosaic genome structure of the Wolbachia *w*Ri strain infecting *Drosophila simulans*. Proc. Nat. Acad. Sci. USA 106, 5725–5730 (2009).

22. Choi, J. Y. & Aquadro, C. F. The coevolutionary period of *Wolbachia pipientis* infecting *Drosophila ananassae* and its impact on the evolution of the host germline stem cell regulating genes. Mol. Biol. Evol. 31, 2457–2471 (2014).

23. Ballard, J. W. O. Comparative genomics of mitochondrial DNA in *Drosophila simulans*. J Mol Evol 51, 64–75 (2000).

24. Ballard, J. W., Hatzidakis, J., Karr, T. L. & Kreitman, M. Reduced variation in *Drosophila simulans* mitochondrial DNA. Genetics 144, 1519–1528 (1996).

25. Ballard, J. W. O. Comparative genomics of mitochondrial DNA in *Drosophila simulans*. J Mol Evol 51, 64–75 (2000).

26. Solignac, M., Vautrin, D. & Rousset, F. Widespread occurence of the proteobacteria Wolbachia and partial cytoplasmic incompatibility in Drosophila melanogaster. C. R. Acad. Sci. 317, 461–479 (1994).

27. Solignac, M. & Monnerot, M. Race formation, speciation, and introgression within *Drosophila simulans, D. mauritiana*, and *D. sechellia* inferred from mitochondrial DNA analysis. Evolution 40, 531–539 (1986).

28. Baba-Aïssa, F., Solignac, M., Dennebouy, N. & David, J. R. Mitochondrial DNA variability in *Drosophila simulans*: quasi absence of polymorphism within each of the three cytoplasmic races. Heredity 61, 419–426 (1988).

29. Montchamp-Moreau, C., Ferveur, J. F. & Jacques, M. Geographic distribution and inheritance of three cytoplasmic incompatibility types in *Drosophila simulans*. Genetics 129, 399–407 (1991).

30. James, A. C. & Ballard, J. W. Expression of cytoplasmic incompatibility in *Drosophila simulans* and its impact on infection frequencies and distribution of *Wolbachia pipientis*. Evolution 54, 1661–1672 (2000).

31. Early, A. M. & Clark, A. G. Monophyly of *Wolbachia pipientis* genomes within *Drosophila melanogaster*: geographic structuring, titre variation and host effects across five populations. Mol. Ecol. 22, 5765–5778 (2013).

32. Richardson, M. F. et al. Population genomics of the Wolbachia endosymbiont in *Drosophila melanogaster*. PLoS Genet. 8, e1003129 (2012).

33. Nunes, M. D. S., Nolte, V. & Schltterer, C. Nonrandom Wolbachia infection status of *Drosophila melanogaster* strains with different mtDNA haplotypes. Mol. Biol. Evol. 25, 2493–2498 (2008).

34. Beckmann, J. F., Ronau, J. A. & Hochstrasser, M. A *Wolbachia* deubiquitylating enzyme induces cytoplasmic incompatibility. Nat Microbiol 2, 17007 (2017).

35. Civetta, A. Rapid Evolution and gene-specific patterns of selection for three genes of spermatogenesis in *Drosophila*. Mol. Biol. Evol. 23, 655–662 (2005).

36. Bauer DuMont, V. L., Flores, H. A., Wright, M. H. & Aquadro, C. F. Recurrent positive selection at *bgcn*, a key determinant of germ line differentiation, does not appear to be driven by simple coevolution with its partner protein *bam*. Mol. Biol. Evol. 24, 182–191 (2006).

37. Flores, H. A., Bubnell, J. E., Aquadro, C. F. & Barbash, D. A. The *Drosophila bag of marbles* gene interacts genetically with Wolbachia and shows female-specific effects of divergence. PLoS Genet. 11, e1005453 (2015).

38. Serbus, L. R. & Sullivan, W. A Cellular basis for Wolbachia recruitment to the host germline. PLoS Pathog 3, e190 (2007).

39. Serbus, L. R., Casper-Lindley, C., Landmann, F. & Sullivan, W. The genetics and cell biology of Wolbachia-host interactions. Annu. Rev. Genet. 42, 683–707 (2008).

40. McGraw, E. A., Merritt, D. J., Droller, J. N. & O'Neill, S. L. Wolbachia density and virulence attenuation after transfer into a novel host. Proc. Nat. Acad. Sci. USA 99, 2918–2923 (2002).

41. Bordenstein, S. R., Marshall, M. L., Fry, A. J., Kim, U. & Wernegreen, J. J. The tripartite associations between Bacteriophage, Wolbachia, and Arthropods. PLoS Pathog 2, e43–10 (2006).

42. Martinez, J. et al. Should symbionts be nice or selfish Antiviral effects of Wolbachia are costly but reproductive parasitism is not. PLoS Pathog 11, e1005021 (2015).

43. Clark, M. E., Veneti, Z., Bourtzis, K. & Karr, T. L. Wolbachia distribution and cytoplasmic incompatibility during sperm development: the cyst as the basic cellular unit of CI expression. Mech. Dev. 120, 185–198 (2003).

44. Chrostek, E., Marialva, M. S. P., Yamada, R., O'Neill, S. L. & Teixeira, L. High anti-viral protection without immune upregulation after interspecies Wolbachia transfer. PLoS ONE 9, e99025 (2014).

45. Chrostek, E. & Teixeira, L. Mutualism breakdown by amplification of Wolbachia genes. PLoS Biol. 13, e1002065 (2015).

46. Chrostek, E. et al. Wolbachia variants induce differential protection to viruses in *Drosophila melanogaster*: A phenotypic and phylogenomic analysis. PLoS Genet. 9, e1003896 (2013).

47. Hoffmann, A. A. et al. Successful establishment of Wolbachia in Aedes populations to suppress dengue transmission. Nature 476, 454–457 (2011).

48. McMeniman, C. J. et al. Host adaptation of a Wolbachia strain after long-term serial passage in mosquito cell lines. App. Environ. Microbiol. 74, 6963–6969 (2008).

49. Osborne, S. E., Iturbe-Ormaetxe, I., Brownlie, J. C., O'Neill, S. L. & Johnson, K. N. Antiviral protection and the importance of Wolbachia density and tissue tropism in *Drosophila simulans*. App. Environ. Microbiol. 78, 6922–6929 (2012).

50. Kondo, N., Shimada, M. & Fukatsu, T. Infection density of Wolbachia endosymbiont affected by co-infection and host genotype. Biol. Lett. 1, 488–491 (2005).

51. Reynolds, K. T., Thomson, L. J. & Hoffmann, A. A. The effects of host age, host nuclear background and temperature on phenotypic effects of the virulent Wolbachia strain popcorn in *Drosophila melanogaster*. Genetics 164, 1027–1034 (2003).

52. Olsen, K., Reynolds, K. T. & Hoffmann, A. A. A field cage test of the effects of the endosymbiont Wolbachia on *Drosophila melanogaster*. Heredity (2001).

53. Boyle, L., O'Neill, S. L., Robertson, H. M. & Karr, T. L. Interspecific and intraspecific horizontal transfer of Wolbachia in *Drosophila*. Science 260, 1796–1799 (1993).

54. Poinsot, D., Bourtzis, K., Markakis, G. & Savakis, C. *Wolbachia* transfer from *Drosophila melanogaster* into *D. simulans*: host effect and cytoplasmic incompatibility relationships. Genetics (1998).

55. McGraw, E. A., Merritt, D. J., Droller, J. N. & O'Neill, S. L. Wolbachia-mediated sperm modification is dependent on the host genotype in *Drosophila*. Proc. R. Soc. B 268, 2565–2570 (2001).

56. Poinsot, D., Montchamp-Moreau, C. & Mercot, H. Wolbachia segregation rate in *Drosophila simulans* naturally bi-infected cytoplasmic lineages. Heredity85 (Pt 2), 191–198 (2000).

57. Boyle, L., O'Neill, S. L., Robertson, H. M. & Karr, T. L. Interspecific and intraspecific horizontal transfer of Wolbachia in *Drosophila*. Science 260, 1796–1799 (1993).

58. Perrot-Minnot & Werren. Wolbachia infection and incompatibility dynamics in experimental selection lines. J. Evol. Biol. 12, 272–282 (1999).

59. Amaya, M., Baranova, A. & van Hoek, M. L. Protein prenylation: A new mode of host-pathogen interaction. Biochemical and Biophysical Research Communications 416, 1–6 (2011).

60. Reinicke, A. T. et al. A *Salmonella typhimurium* effector protein SifA is modified by host cell prenylation and s-acylation machinery. J. Biol. Chem. 280, 14620–14627 (2005).

61. Price, C. T. D., Al-Quadan, T., Santic, M., Jones, S. C. & Abu Kwaik, Y. Exploitation of conserved eukaryotic host cell farnesylation machinery by an F-box effector of *Legionella pneumophila*. J Exp Med 207, 1713–1726 (2010).

62. Signor, S. A., New, F. & Nuzhdin, S. *in review*. An abundance of high frequency variance uncovered in a large panel of Drosophila simulans.

63. Li, H. Aligning sequence reads, clone sequences and assembly contigs with BWA-MEM arXiv. 1–3 (2015).

64. Li, H. et al. The Sequence Alignment/Map format and SAMtools. Bioinformatics 25, 2078–2079 (2009).

65. Meiklejohn, C. D. et al. An Incompatibility between a mitochondrial tRNA and its nuclear-encoded tRNA synthetase compromises development and fitness in *Drosophila*. PLoS Genet. 9, e1003238 (2013).

66. McKenna, A. et al. The Genome Analysis Toolkit: a MapReduce framework for analyzing next-generation DNA sequencing data. Genome Res. 20, 1297–1303 (2010).

67. Danecek, P. et al. The variant call format and VCFtools. Bioinformatics 27, 2156–2158 (2011).

68. Quinlan, A. R. & Hall, I. M. BEDTools: a flexible suite of utilities for comparing genomic features. Bioinformatics 26, 841–842 (2010).

69. Tajima, F. Statistical method for testing the neutral mutation hypothesis by DNA polymorphism. Genetics 123, 585–595 (1989).

70. Ewing, G. & Hermisson, J. MSMS: a coalescent simulation program including recombination, demographic structure and selection at a single locus. Bioinformatics 26, 2064–2065 (2010).

71. Stamatakis, A. RAxML version 8: a tool for phylogenetic analysis and post-analysis of large phylogenies. Bioinformatics 30, 1312–1313 (2014).

72. Izquierdo-Carrasco, F., Smith, S. A. & Stamatakis, A. Algorithms, data structures, and numerics for likelihood-based phylogenetic inference of huge trees. BMC Bioinformatics 12, 470 (2011).

73. Garud, N. R., Messer, P. W., Buzbas, E. O. & Petrov, D. A. Recent selective sweeps in North American *Drosophila melanogaster* show signatures of soft sweeps. PLoS Genet. 11, e1005004 (2015).

74. Ferreira, M. A. R. & Purcell, S. M. A multivariate test of association. Bioinformatics 25, 132–133 (2008).

75. Purcell, S. et al. PLINK: A tool set for whole-genome association and population-based linkage analyses. Am. J. Hum. Genet. 81, 559–575 (2007). 76.

76. FlyBase Curators, Swiss-Prot Project Members InterPro Project Members. Gene Ontology annotation in FlyBase through association of InterPro records with GO terms. (2004).

77. Babu, K., Bahri, S., Alphey, L. & Chia, W. Bifocal and PP1 interaction regulates targeting of the R-cell growth cone in *Drosophila*. Dev. biol. 288, 372–386 (2005).

78. Bennett, D., Szoor, B., Gross, S., Vereshchagina, N. & Alphey, L. Ectopic expression of inhibitors of Protein phosphatase type 1 (PP1) can be used to analyze roles of PP1 in *Drosophila* development. Genetics 164, 235–245 (2003).

79. Gaudet, P., Livstone, M. & Thomas, P. Gene Ontology annotation inferences using phylogenetic trees. GO Reference Genome Project. (2010).

80. Giagtzoglou, N. et al. *dEHBP1* controls exocytosis and recycling of *Delta* during asymmetric divisions. J. Cell Biol. 196, 65–83 (2012).

81. Cronin, S. J. F. et al. Genome-wide RNAi screen identifies genes involved in intestinal pathogenic bacterial infection. Science 325, 340–343 (2009).

82. Newton, I. L. G., Savytskyy, O. & Sheehan, K. B. Wolbachia utilize host actin for efficient maternal transmission in *Drosophila melanogaster*. PLoS Pathog 11, e1004798 (2015).

83. Chiba, A. Early development of the *Drosophila* neuromuscular junction: a model for studying neuronal networks in development. Int. Rev. Neurobiol. (1999).

84. Yampolsky, L. Y. & Stoltzfus, A. The exchangeability of amino acids in proteins. Genetics 170, 1459–1472 (2005).

85. Creixell, P., Schoof, E. M., Tan, C. S. H. & Linding, R. Mutational properties of amino acid residues: implications for evolvability of phosphorylatable residues. Philos. Trans. R. Soc. B 367, 2584–2593 (2012).

86. Leone, A. et al. Evidence for nm23 RNA overexpression, DNA amplification and mutation in aggressive childhood neuroblastomas. Oncogene 8, 855–865 (1993).

87. Ishiko, A. et al. A novel leucine to valine mutation in residue 7 of the helix initiation motif of Keratin10 leads to bullous congenital ichthyosiform erythroderma. J. Inv. Dermatol. 116, 991–992 (2001).

88. Mahowald, A. P. Assembly of the *Drosophila* germ plasm. Int. Rev. Cytol. 203, 187–213 (2001).

89. Kose, H. & Karr, T. L. Organization of *Wolbachia pipientis* in the *Drosophila* fertilized egg and embryo revealed by an anti-Wolbachia monoclonal antibody. Mech. Dev. 51, 275–288 (1995).

90. Serbus, L. R. et al. A feedback loop between Wolbachia and the *Drosophila gurken* mRNP complex influences Wolbachia titer. J. Cell Sci. 124, 4299–4308 (2012).

91. Ballard, J. W. O. Comparative genomics of mitochondrial DNA in *Drosophila simulans.* J. Mol. Evol. 51, 64–75 (2000).

92. Vinh, D. B. N., Ko, D. C., Rachubinski, R. A., Aitchison, J. D. & Miller, S. I. Expression of the *Salmonella spp.* virulence factor SifA in yeast alters Rho1 activity on peroxisomes. Mol. Biol. Cell 21, 3567–3577 (2010).

93. Brumell, J. H., Goosney, D. L. & Finlay, B. B. SifA, a type III secreted effector of *Salmonella typhimurium*, directs *Salmonella*-induced filament (Sif) formation along microtubules. Traffic 3, 407–415 (2002).

94. Siozios, S. et al. The diversity and evolution of Wolbachia ankyrin repeat domain genes. PLoS ONE 8, e55390 (2013).

95. Bork, P. Hundreds of ankyrin-like repeats in functionally diverse proteins: mobile modules that cross phyla horizontally Proteins 17, 363–374 (1993).

96. Habyarimana, F. et al. Role for the Ankyrin eukaryotic-like genes of *Legionella pneumophila* in parasitism of protozoan hosts and human macrophages. Environ. Microbiol. 10, 1460–1474 (2008).

97. Veneti, Z., Clark, M. E., Karr, T. L., Savakis, C. & Bourtzis, K. Heads or Tails: Host-Parasite Interactions in the *Drosophila*-Wolbachia System. App. Env. Microbiol. 70, 5366–5372 (2004).

